# Physiological and transcriptomic response to methyl-coenzyme M reductase limitation in *Methanosarcina acetivorans*

**DOI:** 10.1101/2023.12.13.571449

**Authors:** Grayson L. Chadwick, Gavin A. Dury, Dipti D. Nayak

## Abstract

Methyl-coenzyme M reductase (MCR) catalyzes the final step of methanogenesis, the microbial metabolism responsible for nearly all biological methane emissions to the atmosphere. Decades of biochemical and structural studies have generated detailed insights into MCR function *in vitro*, yet very little is known about the interplay between MCR and methanogen physiology. For instance, while it is routinely stated that MCR catalyzes the rate-limiting step of methanogenesis, this statement has not been categorically tested. Here, to gain a more direct understanding of MCR’s control on the growth of *Methanosarcina acetivorans,* we generate a strain with an inducible *mcr* operon on the chromosome, allowing for careful control of MCR expression. We show that MCR is not growth rate limiting in substrate-replete batch cultures. However, through careful titration of MCR expression, growth-limiting state(s) can be obtained. Transcriptomic analysis of *M. acetivorans* experiencing MCR-limitation reveals a global response with hundreds of differentially expressed genes across diverse functional categories. Notably, MCR limitation leads to a strong induction of methylsulfide methyltransferases, likely due to insufficient recycling of metabolic intermediates. In addition, the *mcr* operon does not seem to be transcriptionally regulated, i.e., it is constitutively expressed, suggesting that the overabundance of MCR might be beneficial when cells experience nutrient limitation or stressful conditions. Altogether, we show that there is wide range of cellular MCR concentrations that can sustain optimal growth, suggesting that other factors like anabolic reactions might be rate-limiting for methanogenic growth.

**Importance:** Methane is a potent greenhouse gas that has contributed to *ca.* 25% of global warming in the post-industrial era. Atmospheric methane is primarily of biogenic origin, mostly produced by microorganisms called methanogens. In methanogens, methyl-coenzyme M reductase (MCR) catalyzes methane formation. Even though MCR comprises *ca.* 10% of the cellular proteome, it is hypothesized to be growth-limiting during methanogenesis. Here, we show that *Methanosarcina acetivorans* grown under standard laboratory conditions produces more MCR than its cellular demand for optimal growth. The tools outlined in this study can be used to refine metabolic models of methanogenesis and assay lesions in MCR in a higher throughput manner than isolation and biochemical characterization of pure protein.

## Introduction

Methyl coenzyme M reductase (MCR) catalyzes the final step of methane production in methanogenic archaea (1). The active enzyme consists of three subunits in an α_2_β_2_γ_2_ stoichiometry that is present in very high abundance in the cytosol. The operon encoding MCR dominates methanogen transcriptomes, where it is invariably found to be one of the most abundant mRNAs (2), and MCR accounts for roughly 10% of cytoplasmic proteome (1, 3). MCR is also the first step of anaerobic methane oxidation in anaerobic methanotrophic archaea, where it is found in similarly high levels in transcriptomes (4, 5) and proteomes (6, 7). Despite its abundance, the isolation and biochemical characterization of active MCR is challenging. The active site contains a nickel porphyrin cofactor (F430) which is exquisitely oxygen sensitive, and even when the enzyme is purified in the absence of oxygen it can enter inactive product-inhibited states. While a few protocols have been developed to purify the active enzyme, or re-activate inactive states, from *Methanothermobacter marburgensis*, these are not broadly applicable to other methanogens, especially genetically tractable strains like *Methanococcus maripaludis* or *Methanosarcina spp.* (1). Based on these challenges, even straightforward experiments to measure kinetic parameters like substrate affinities (K_m_) or turnover rate (k_cat_) have taken decades of effort from multiple research groups and have not been expanded to cover any significant phylogenetic diversity.

In contrast to the advances made in understanding the biochemical properties of the *M. marburgensis* MCR in isolation, holistic questions that address the biogenesis and function of MCR, likely mediated by interaction with other proteins, as well as with methanogen physiology more broadly, have received significantly less attention. We know very little about the proteins involved in the assembly, activation, and degradation of MCR *in vivo.* Similarly, regulatory processes that control the expression of MCR in response to environmental cues like resource availability or stress response have barely been studied. The only system where regulation has been investigated is the differential expression of the two isoforms of MCR present in *Methanothermobacter spp*.(8), however most methanogens carry only a single copy of MCR (1). How, and to what extent, methanogens control the amount of MCR and its activity when conditions are unfavorable is an understudied but important question due to the growing interest in inhibiting methanogenesis for the purpose of reducing methane emissions, as well as using methanogens as chassis for bioengineering wherein major metabolic end products besides methane are desired (9).

Recent advances in genetic tools available for the study of methanogens have facilitated targeted mutagenesis of genes encoding hypothesized accessory proteins of MCR and provided an avenue to investigate their cellular functions. Using this approach, the identity and function of many MCR-associated proteins involved in the installation of post-translational modifications and insertion of F430 have been successfully studied(10–12). One surprising outcome of these genetic studies is that the loss of highly conserved MCR-associated proteins often have little to no significant effect on cell growth, even though MCR is essential in all methanogenic archaea to the best of our knowledge (10–12). A possible explanation for these results is that MCR is not rate-limiting for growth under standard laboratory conditions. Therefore, even mutations that result in a substantial detrimental impact to MCR function may be tolerated without a notable growth defect, as has been recently suggested (1).

No direct evidence linking MCR abundance to growth and methanogenesis is currently available, and there are differing views literature. A recent study used a kinetic and stoichiometric hybrid approach to model the growth of *Methanosarcina barkeri* on methanol (13). Here, they showed MCR has a high control coefficient (of ∼0.9) during growth on high methanol concentrations (> 15 mM) i.e., that small changes in MCR levels have a dramatic impact on growth rate under typical batch-culture conditions. This model as well an older kinetic model for *Methanosarcina acetivorans* (14) corroborate a long-standing hypothesis that MCR is a rate-limiting enzyme during methanogenesis (8, 15). In contrast, studies that have targeted F430 biosynthesis, either through the omission of nickel in the growth medium or through the addition of levulinic acid, which inhibits porphyrin biosynthesis (3, 16, 17). These studies show that a modest decrease in F430 abundance has no effect on cell growth and only a drastic reduction in F430 levels, by 5-fold or more, leads to growth defects and can alter subcellular localization of MCR. Taken together, these studies are consistent with the notion that MCR is present in excess in nickel replete medium typically used for laboratory cultivation of methanogens (3). One caveat of these studies is that nickel and porphyrins are present in many other bioenergetic enzymes essential for methanogenesis hence it is difficult to attribute any observed growth phenotypes entirely to MCR. Additionally, carbon monoxide supplementation inhibits methanogenesis in *Methanosarcina acetivorans* and growth under these conditions produces large quantities of acetate, formate, and methylsulfides and only a small amount of methane (18–20). Acetogenic growth of *M. acetivorans* can be further amplified in mutants where methane production is nearly completely abolished (21). While it has been suggested that carbon monoxide is partially inhibitory to MCR (18), it is not clear that there is biochemical basis for this notion. Notably, even under conditions where methane production is an insignificant portion of catabolism, MCR remains essential and cannot be deleted (21).

Based on the current literature it is unclear if MCR is present in excess during laboratory growth, and if so, why such a substantial portion of their proteome and transcriptome might be allocated to it - especially when there is well established precedent that methanogens have elaborate mechanisms for modifying the expression of other metabolic genes (22). Here, to better understand this interplay between MCR abundance and cell growth, we investigated the physiology of the genetically tractable strain *M. acetivorans* carrying an inducible MCR operon. Our results clearly demonstrate that wild-type expression of MCR far exceeds the cellular demand, but that MCR-limited growth is indeed possible at significantly lower levels of expression. Under these MCR-limiting conditions there is a global transcriptional shift that alters the expression of hundreds of genes involved in a variety of cellular processes beyond methanogenesis.

## Results and Discussion

### Halting the transcription of the MCR operon leads to linear growth

*M. acetivorans* strain WWM60 encodes a *tetR* gene under the control of the *mcrB* promoter from *Methanosarcina barkeri* Fusaro in place of the *hpt* locus (MA0717 or MA_RS03755), enabling tetracycline-based control of gene expression and protein production (23). We used WWM60 as the genetic background to introduce a *tetO1* operator site in the promoter of the *mcr* operon and generated the mutant strain DDN032, wherein the expression of the *mcr* operon can be titrated by the addition of tetracycline to the growth medium (**Fig. 1A**). Whole genome resequencing of DDN032 verified the desired chromosomal change as well as the absence of any suppressor mutation(s) or off-target effects due to CRISPR editing (**Fig. S1**). Unless specified, this strain was passaged in media containing 100 µg/ml tetracycline, which is considered full induction of the P*mcrB*(*tetO1*) promoter (23).

**FIG 1:**
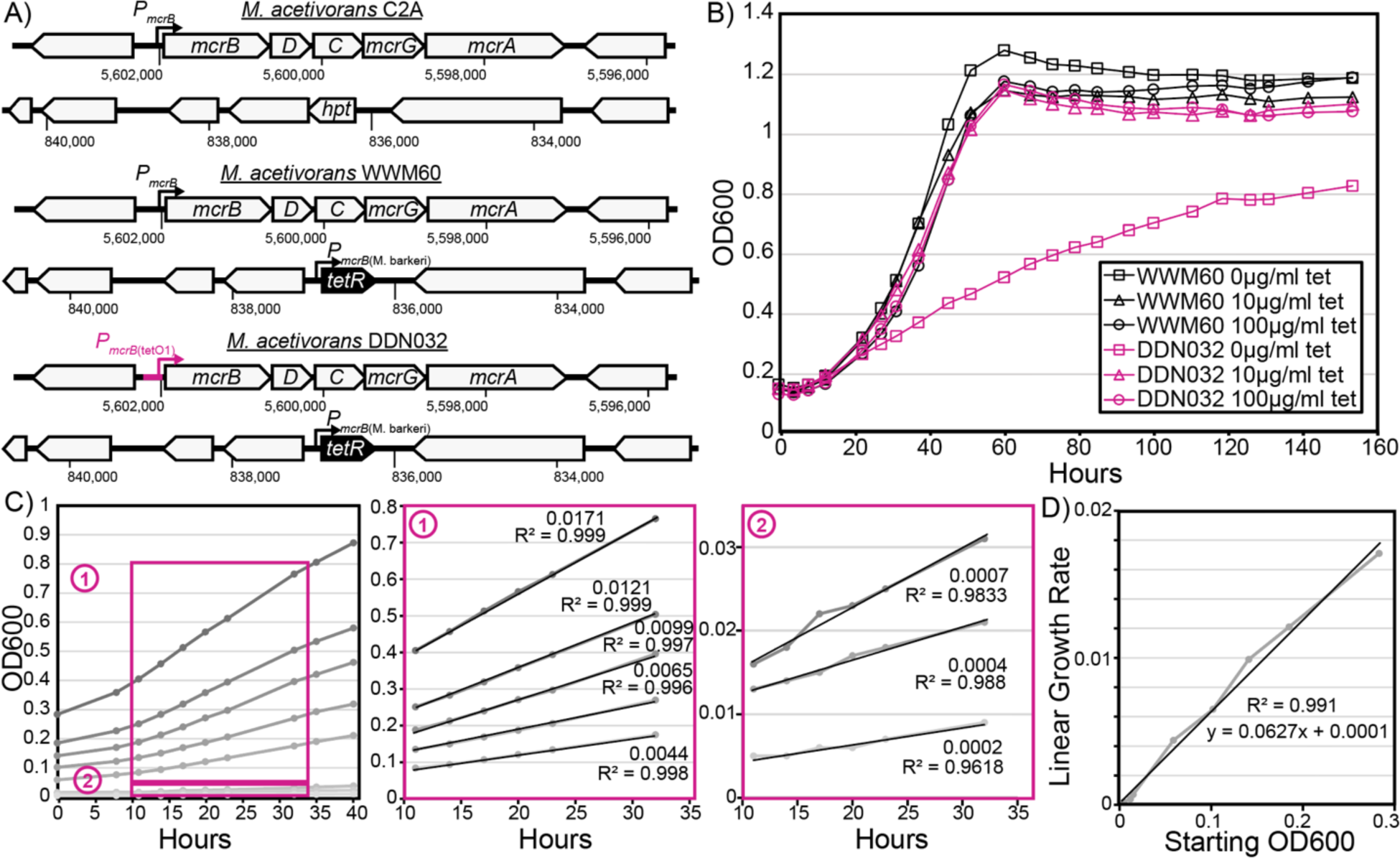
Mutant construction and growth characteristics. (A) Genotype of wild-type *M. acetivorans* C2A, the strain capable of inducible expression via the chromosomal integration of *tetR* (WWM60) and the inducible MCR strain generated in this study (DDN032). (B) Growth characterization of DDN032 on high-salt trimethylamine media demonstrates a linear growth phenotype upon transfer into media lacking tetracycline (representative growth curves shown). (C) Linear growth curves of DDN032 in tetracycline-free media at various starting optical densities. Boxes 1 and 2 show details of regions used for linear regression calculations. Slopes and R^2^ values are shown beside each line in boxes 1 and 2. (D) The slopes of the linear regressions from panel C are plotted as a function of the starting optical density demonstrating a strong linear relationship.

To determine the effect of MCR expression on growth, we inoculated exponential phase cultures of WWM60 and DDN032 (previously grown in medium with 100 µg/ml tetracycline and washed in tetracycline-free media three times) in media containing 0, 10, or 100 µg/ml tetracycline with trimethylamine as their carbon source. DDN032 grew indistinguishably from WWM60 in medium supplemented with 10 or 100 µg/ml tetracycline, however, there was a clear distinction between the two strains in the absence of tetracycline (**Fig. 1B**). Under these conditions, DDN032 has a biphasic growth curve starting with a short phase of exponential growth (corresponding to approximately one doubling) followed by linear growth. Linear growth can result from the dilution of a growth-limiting substance upon cell division such that the two daughter cells will have half the growth rate of the parental cell (24). We hypothesize that the linear growth observed here is due to continuous dilution of growth-limiting amounts of MCR protein once MCR production halts. A short period of exponential growth before onset of linear growth has previously been observed for nickel limitation in *Methanothermobacter thermautotrophicus* (3).

If MCR limitation is leading to linear growth of DDN032, then based on prior art (24), the rate of linear growth would be directly proportional to the amount of MCR (the limiting substance) in the inoculum. To test this hypothesis, cultures of DDN032 grown with 100 µg/ml tetracycline were triply washed and inoculated into tubes without tetracycline at varying starting optical densities (**Fig. 1C**). Within 10 hours, cultures entered linear growth in which the rate of increase in optical density was entirely dependent on the starting optical density, consistent with the prediction of MCR being the limiting resource giving rise to linear growth (**Fig. 1D**). We were able to observe slow linear growth for weeks in cultures seeded with a low inoculum, however, after prolonged of incubation, escape mutations were commonly observed due to the inactivation of *tetR* by endogenous transposon insertion (**Fig. S2**).

### Measuring the expression threshold for aberrant growth due to MCR limitation

With clear evidence that complete MCR suppression leads to aberrant growth in *M. acetivorans*, we sought to determine the minimum amount of tetracycline necessary for wild-type levels of growth. To determine this induction threshold, DDN032 was inoculated into media supplemented with a series of tetracycline concentrations. In two separate experiments, growth was analyzed in quintuplicate cultures with a wide range of tetracycline concentrations (Experiment 1) or quadruplicate cultures with a fine range of tetracycline concentrations (Experiment 2), all with methanol as the carbon source (**Fig. 2A**, **S3-4**). For quantitative comparisons between all growth conditions, we calculated growth rates based on exponential fits of the initial 1-2 doublings (**Fig. 2B**). In both experiments, when growth rates of DDN032 were compared to WWM60 there was no significant difference in growth at tetracycline concentrations ≥2.5µg/ml, whereas a detectable growth defect was observed at tetracycline concentrations ≤1.75 µg/ml in Experiment 1 and ≤2 µg/ml in Experiment 2.

**FIG 2:**
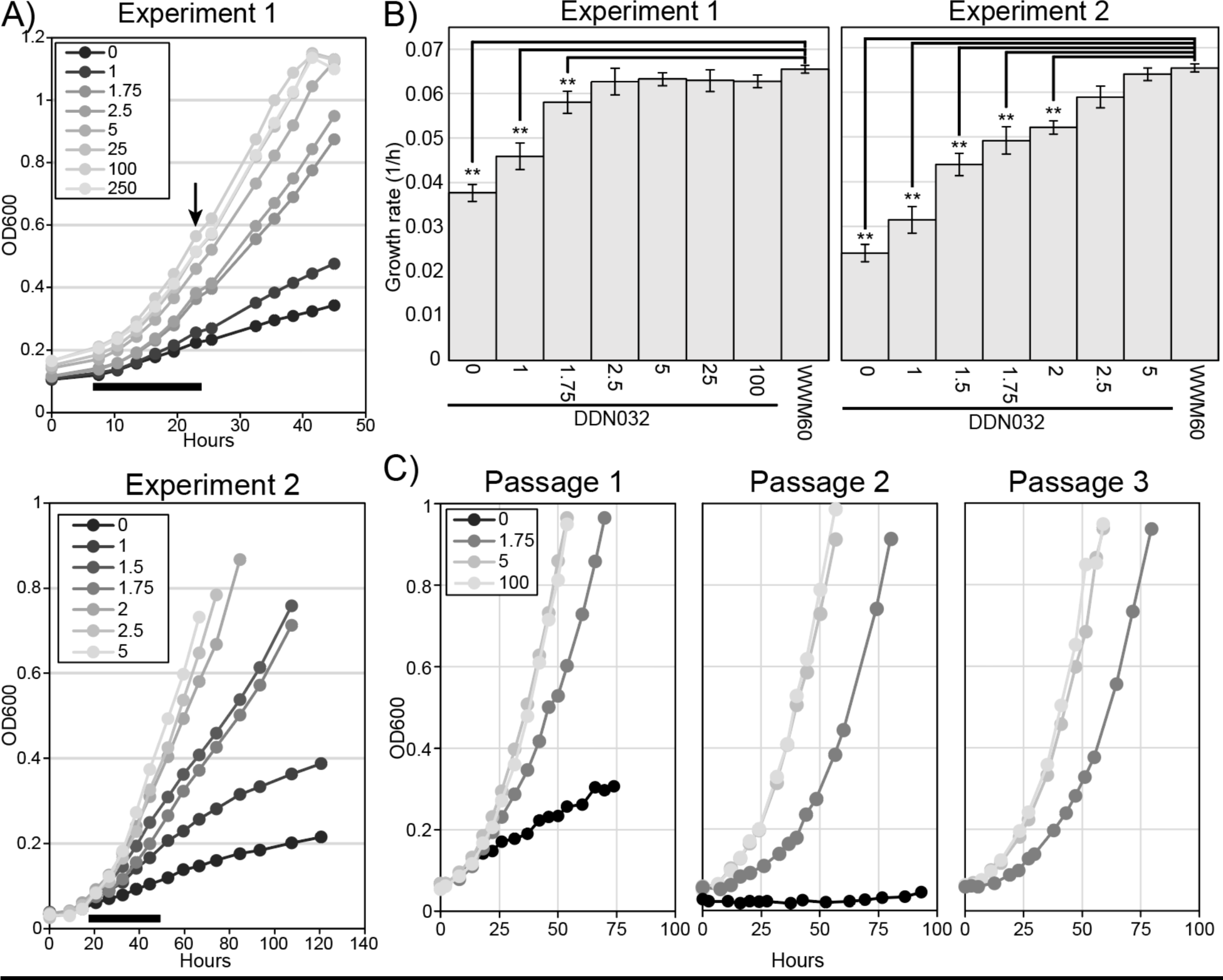
Effect of tetracycline on growth rate. (A) Cultures of DDN032 pregrown in media with 100µg/ml tetracycline were triply washed an inoculated into Balch tubes with 8 different concentrations of tetracycline (µg/ml) in quintuplicate (Experiment 1, top) and 7 different concentrations in quadruplicate (Experiment 2, bottom). Representative growth curves are shown here; all growth data can be found in **Fig. S3-4**. Black bar below growth curves shows the time range where growth data was calculated for panel B, and the arrow indicates in Experiment 1 where three of five replicates were sacrificed for RNA extraction (see **Fig. 3-4**). (B) Growth rates for DDN032 and WWM60 in the conditions indicated. Error bars indicate standard deviations of quintuplicate cultures for Experiment 1 and quadruplicate for Experiment 2. Growth rates were calculated from exponential fits in the region shown by a black bar in panel A as discussed in the text. Note: growth for WWM60 with 100µg/ml tetracycline was only carried out in Experiment 1. Growth rates that are significantly different than WWM60 with 100µg/ml tetracycline ANOVA and Tukey’s Honest Significant Difference test are indicated (** p-value ≤ 0.01). (C) Sequential passaging of DDN032 at 1.75µg/ml tetracycline reveals reproducibly slower exponential growth (also see **Fig S5**).

For the lowest tetracycline concentrations considered above, there is a clear transition into linear growth, so the exponential growth rates reported in **Fig 2B** do not correspond to a sustainable growth phenotype. However, it is possible that just below the apparent 2.5µg/ml threshold, DDN032 may achieve sustained exponential growth at lower-than-wild-type levels. To assess this, we carried out three passages of DDN032 cultures at various tetracycline concentrations including 1.75 µg/ml (**Fig. 2C**). Even after three passages, exponential growth was observed in cultures with 1.75 µg/ml to be slower than at 5 and 100 µg/ml tetracycline, suggesting that it is possible for true exponential growth to be achieved in an MCR-limited manner (**Fig. 2C, Fig S5**).

### Effect of tetracycline on transcriptome and MCR protein abundance

To quantify MCR repression and the resulting transcriptomic response, we extracted and sequenced total RNA from triplicate cultures grown at various tetracycline concentrations at the timepoint shown in **Fig. 2A**. This timepoint was chosen as it represented one of the earliest points where the growth curves of the conditions with different concentrations of tetracycline started diverging (**Fig. 2A**). Transcription of the *mcrA* gene in DDN032 decreased continuously from an FPKM value of ∼850 at 100 µg/ml tetracycline to ∼6 at 0 µg/ml (**Fig. 3A**). Notably, even though no growth difference was observed between WWM60 and DDN032 supplemented with 100 µg/ml tetracycline, there is approximately half the level of *mcrA* transcription in the latter. This inability to reach full induction is possibly because the *tetO1* operator decrease the expression strength of the P*mcrB*(*tetO1*) promoter relative to the native promoter. When plotted against *mcrA* transcript abundance, the growth rates are indistinguishable from the parental strain until transcription drops to approximately one third of the parental level, suggesting a large range of MCR-replete growth (**Fig. 3B**). This pattern also holds true at the protein level where the abundance of MCR was measured with polyclonal antibodies raised against *M. acetivorans* McrA and revealed a clear decrease in MCR abundance at different concentrations of tetracycline for DDN032 (**Fig. 3C**).

**FIG 3:**
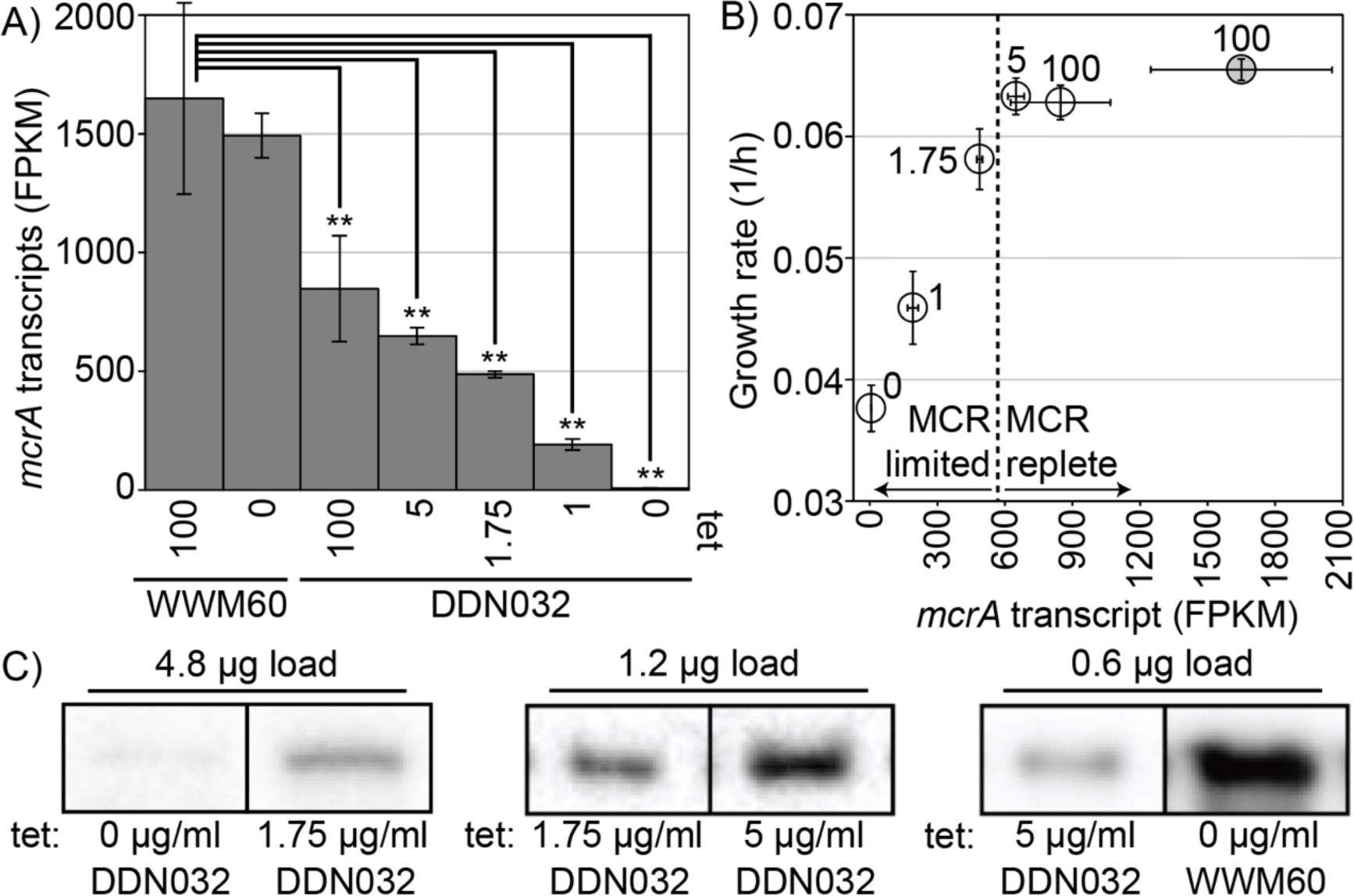
MCR transcript and protein response to tetracycline concentration. Cultures grown at 0, 1, 1.75, 5 or 100 µg/ml tetracycline (DDN032) or 0 and 100 µg/ml (WWM60) were used for RNA sequencing. (A) Transcriptional response of the *mcrA* gene to tetracycline concentration in WWM60 and DDN032. Transcript abundances that are significantly different than WWM60 with 100µg/ml tetracycline by ANOVA and Tukey’s Honest Significant Difference test are indicated (** p-value ≤ 0.01). Error bars represent standard deviations of biological triplicates. (B) Grow rates of cultures vs. the *mcrA* transcript level, open circles DDN032, shaded circle WWM60 with 100 µg/ml tetracycline. Error bars represent standard deviations of biological triplicates, dashed line represents an apparent switch between MCR replete and MCR limited growth and numbers next to the points indicate tetracycline concentration (µg/ml). (C) Representative bands from western blot analysis with antibodies raised against McrA from *Methanosarcina acetivorans*. Comparisons between MCR abundance at different levels of tetracycline concentrations (listed below each band) should only be made with bands resulting from the same total protein load (listed above each band). Bands with the same total protein load are noncontiguous selections from the same image; full blots showing relative concentrations by dilution to extinction are available (**Figure S6**). Note: for DDN032 grown with 1µg/ml only two replicates were available for RNA sequencing.

At the global level there is little difference between the transcriptional profile of WWM60 grown with or without tetracycline. Only a single gene was found to be significantly differentially expressed between the two WWM60 conditions, a putative ABC transporter substrate binding protein (MA0887), indicating that tetracycline alone has little effect on the *M. acetivorans* transcriptome (**Fig. 4A-B, Supplementary Tables 4-5)**. However, the transcriptomic profile of DDN032 was substantially different than that of WWM60 at all tetracycline concentrations investigated (**Fig. 4A**), and the number of genes differentially expressed between DDN032 and WWM60 increased as tetracycline concentration decreased (**Fig. 4B**). Principal component analysis revealed the overall transcriptome structure follows a similar trend as the *mcrA* gene itself (**Fig. 4A**). Principle component 1, which accounts for 63% of the variance in the transcriptome, falls along a gradient of MCR expression. As expected, the most significantly downregulated genes at 1 and 0 µg/ml tetracycline belong to the *mcr* operon (**Fig. 4B**). A few other important genes follow the same trend. The expression of *cfbE* (MA3630), the F_430_ synthetase (25), seems to match that of the *mcr* operon, suggesting the existence of a feedback mechanism to reduce F_430_ biosynthesis in response to MCR limitation. Similarly, a gradual decrease can be observed in *mmp10* (MA4551) which encodes the protein responsible for the posttranslational modification of a conserved arginine residue in MCR (26, 27) (**Fig. 4C**). Catabolic genes that are highly expressed under normal conditions such as the methanol specific methyltransferase isoform 1 *mtaCB1* (MA0455-MA0456) and genes of the MTR complex, are also significantly decreased in expression upon MCR limitation.

**FIG 4:**
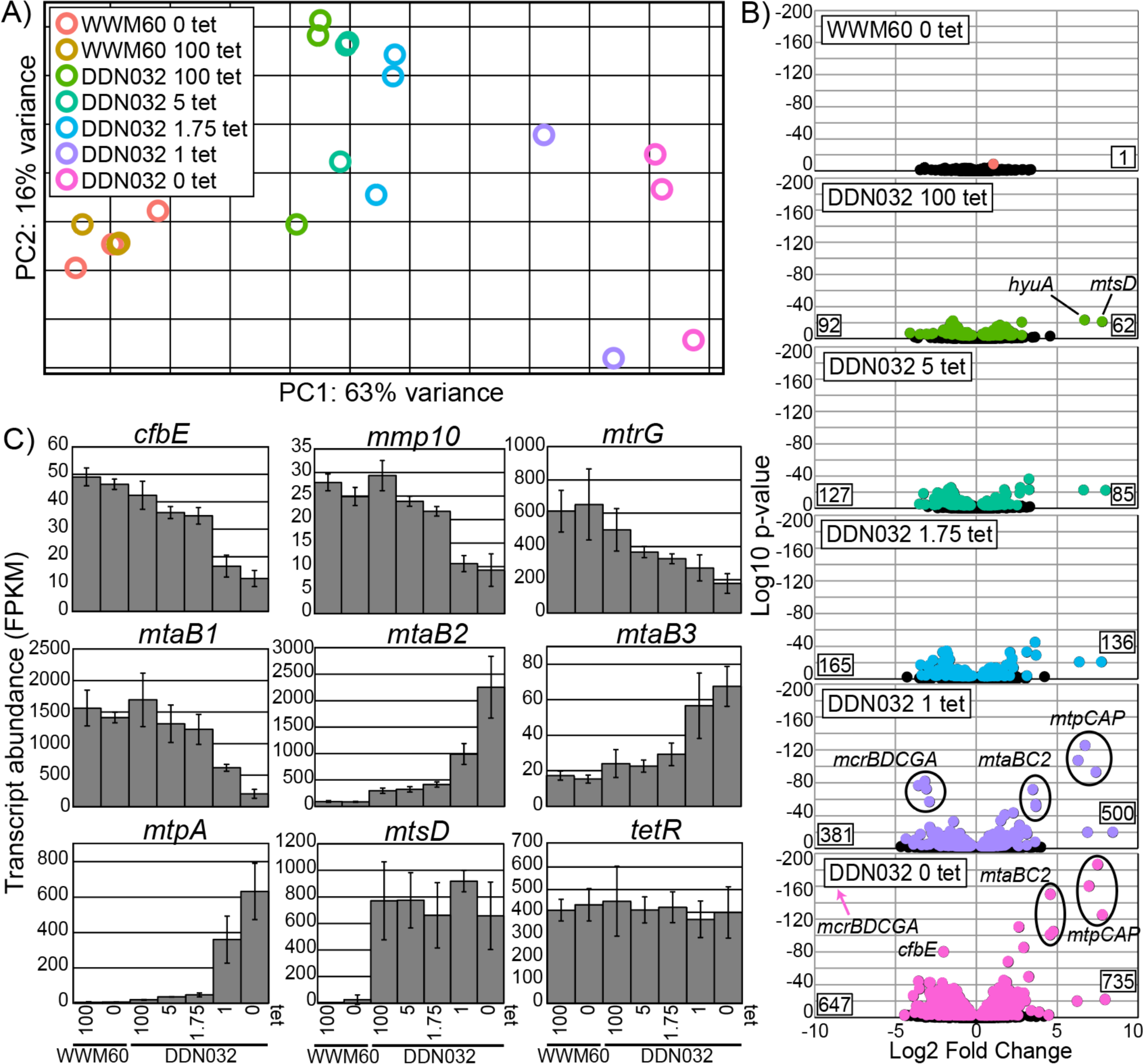
Global transcriptomic response to MCR limitation. (A) Global transcriptomic response to different tetracycline concentrations captured by principal component analysis. (B) Volcano plots depicting the log2-fold change for genes of all conditions compared to WWM60 grown on 100 µg/ml tetracycline. Negative values indicate lower expression relative to WWM60 on 100 µg/ml. All genes significantly differentially expressed based on multiple comparison adjusted P-value cutoff of 0.01 are shown in color, non-significant genes in black. Numbers on the bottom left and right corners of each plot represent the total number of genes significantly down or upregulated, respectively. Certain genes exhibiting strong fold change differences mentioned in the text are highlighted. (C) Specific gene expression profiles as a function of tetracycline concentration. Error bars represent standard deviations of biological triplicates. For *mtpA* and *mtsD* all DDN032 conditions are significantly upregulated compared to WWM60 at a multiple comparison corrected P-value of <0.01 as determined by DESeq2. *tetR* is not significantly differentially expressed under any conditions. Note: for DDN032 grown with 1µg/ml only two replicates were available for RNA sequencing.

Many genes also respond positively to MCR limitation. The *mtaCB2* (MA4391-MA4392) genes that encode the methanol methyltransferase isoform 2 are typically expressed in late-exponential or stationary phase. However, during MCR limitation, and the concomitant decrease in *mtaCB1* expression, these genes are strongly upregulated even in early-mid exponential phase (**Fig. 4B, C**). Methanol methyltransferase isoform 3 encoded by *mtaBC3* (MA1616-MA1617) follows a similar trend, albeit to a lesser degree. By far the most dramatic increase in expression occurs in two of the methyl-sulfide specific methyltransferases, *mtpCAP* (MA4164-MA4166) and *mtsD* (MA0859), which increase in expression approximately 100-fold (**Fig. 4B, C**). Notably, the gene encoding *tetR* does not significantly change in expression level under any of our experimental conditions. This is particularly interesting given that *tetR* is driven by a p*mcrB* promoter and suggests that the *mcr* operon might be constitutively expressed in the cell.

## Conclusions

Here we have provided the first investigation into the physiological and transcriptomic response of a methanogenic archaeon specifically to MCR limitation. We find that MCR is not limiting *M. acetivorans* growth under standard laboratory conditions, and this observation may explain why some universally conserved MCR-associated proteins can be deleted with little to no effect on growth in this system (10–12). Decreasing MCR can lead to two forms of growth limitation, linear growth under extreme MCR limitation, or slower, sustained exponential growth under less drastic limitation (**Fig. 2)**. While this growth-limiting state cannot be maintained indefinitely due to the accumulation of escape mutations, we anticipate that the threshold tetracycline concentration at which a growth defect occurs may be a useful diagnostic feature in assessing the relative fitness of MCR mutants or strains lacking certain conserved MCR-associated proteins. This approach, particularly if coupled with high throughput growth assays, may enable the screening and initial characterization of many MCR variants, far more than is feasible through existing biochemical approaches.

The global transcriptional response to MCR limitation shared similarities with prior observations made of *M. acetivorans* under a variety of stressful conditions. In particular, the upregulation of the *mtpCAP* operon and *mtsD* is reminiscent of increased methylsulfide production during growth on carbon monoxide (18, 19, 28), and in the absence of HdrABC (29) or pyrrolysine (30). In some of these cases, it has been hypothesized that limitation in MCR activity or change in electron flow results in a buildup of methyl-coenzyme M which can then be relieved by an increase in expression of methyl-sulfide methyltransferases, possibly to help maintain redox balance. While there is no evidence for how this alternate pathway might allow *M. acetivorans* to conserve energy, the results presented here are consistent with the idea that methyl-coenzyme M build-up may induce these alternative methyl transferase systems. It is interesting to note in this context that the MTR complex is significantly downregulated, suggesting that if methyl-coenzyme M build-up is indeed occurring, that the oxidative branch of the methylotrophic pathway is not a viable outlet for these methyl groups, presumably due to a build-up of electron carriers. It is also interesting that while MCR itself may not be actively regulated, as evidenced by the lack of expression change of *tetR*, MCR-associated proteins such as *mmp10* and *cfbE* are down-regulated, suggesting a feed-back to the expression of MCR-associated proteins. While this trend is not universally true (e.g. *ycaO*, the McrA-glycine thioamidation protein is not significantly regulated), the list of differentially expressed genes presented here may lead to the discovery of additional MCR-related systems of unknown function.

Altogether, we have developed a genetic platform to conclusively demonstrate that MCR does not mediate the rate-limiting step in *M. acetivorans* during routine laboratory growth conditions. While these data are consistent with prior observations from studies with *Methanothermobacter spp.*, they diverge from predictions made by metabolic models of *Methanosarcina spp.* during methanol growth(13). Clearly, more physiological studies like ours are required to bridge the gap between research with enzymes in isolation and systems-level analyses of methanogens. While this tool in and of itself will prove to be especially useful to study the properties of MCR mutants and of mutations in MCR-associated proteins, this experimental framework can be expanded to other important enzymes like HdrDE, HdrABC and Mtr to ultimately obtain a robust and quantitative view of methanogenesis.

## Materials and Methods

### CRISPR-editing plasmid construction and mutant generation

A target sequence (GTGGACACTTAAAAACGACG) for the *mcrB* promoter in *M. acetivorans* was identified using the CRISPR site finder tool in Geneious Prime version 11.0 with the following parameters: a) an NGG protospacer adjacent motif (PAM) site at the 3’ end and b) no off-target matches allowed. A single guide RNA (sgRNA) was synthesized as a gblock gene fragment from Integrated DNA technologies (Coralville, IA, USA) using the target sequence. The sgRNA and a homology repair template to insert the *tetO1* operator site in the promoter of the *mcrBDCGA* operon were cloned into the Cas9 containing vector pDN201 as described previously (31) to generate pGLC001. pGLC001 was digested with PmeI and a repair template introduced which included the P*mcrB*(*tetO1*) promoter in place of the native *pMcrB* sequence generating pGLC002. The sequences of pGLC001 and pGLC002 were verified by Sanger sequencing at the Barker sequencing facility at University of California, Berkeley. A cointegrate of pGLC002 and pAMG40 was generated using the Gateway BP Clonase II Enzyme mix per the manufacturer’s instructions (Thermo Fisher Scientific, Waltham, MA, USA) and named pGLC003. All *E. coli* transformations were conducted with WM4489 (32) as described previously.

A 10 mL culture of *M. acetivorans* in high salt (HS) medium with 50 mM trimethylamine (TMA) in late-exponential phase was used for liposome-mediated transformation with pGLC003 as described previously (33). Transformants were plated in agar solidified HS medium with 50 mM TMA, 100 µg/ml tetracycline, and 2 µg/ml puromycin and incubated in an intra-chamber incubator at 37 °C with H_2_S/CO_2_/N_2_ (1000 ppm/20%/balance) in the headspace. Colonies were screened for the mutation and sequence verified by Sanger sequencing at the Barker sequencing facility at University of California, Berkeley. Several colonies that tested positive for the desired mutation were streaked out on HS medium with 50 mM TMA, 100 µg/ml tetracycline, 20 µg/ml 8ADP to cure the mutagenic plasmid. Plasmid cured mutants were verified by screening for the absence of the *pac* gene present on the plasmid with PCR. A single isolate of the plasmid cured mutant was grown in liquid culture with 50 mM TMA and 100 µg/ml tetracycline and saved as DDN032. All primers, plasmids, and strains used in this study are listed in Supplementary Tables 1-3, respectively.

### Growth measurements

All growth experiments were conducted using either WWM60 (*M. acetivorans* Δ*hpt*:: P*mcrB-tetR*) (23) or DDN032 (WWM60-P*mcrB*(*tetO1*)-*mcrBDCGA*), a strain with the chromosomal *mcr* genes containing a *tetO1* operator site inserted in the promoter. All growth analyses were in 10 ml of high salt (HS) media containing methanol (125mM) or TMA (50mM) as a carbon source and a pressurized CO_2_/N_2_ (20:80) headspace as previously described (34). Various concentrations of tetracycline were added to the media as indicated, requiring the media to be protected from light to prevent degradation. Anaerobic tetracycline stocks were made fresh in anaerobic water on the day of the inoculation as described previously (23).

All *M. acetivorans* doubling times were determined by measuring the optical density (at 600 nm) of cultures grown in Balch tubes containing 10 ml HS media with media additions as indicated. All optical density measurements were made using a UV-Vis spectrophotometer (Gensys 50, Thermo Fisher Scientific). Doubling times were determined using the best fit line of the log_2_ transformed optical density data with maximal R^2^ values.

### DNA extraction and sequencing

Genomic DNA was extracted from a 10 ml late-exponential phase culture of DDN032 in HS medium with 125 mM methanol and 100 µg/ml tetracycline as well as the escape mutant in HS medium with 125 mM methanol using a Qiagen blood and tissue kit per the manufacturer’s instructions (Qiagen, Hilden Germany). Library preparation and Illumina sequencing (150 bp paired end reads) was conducted at Seqcenter (Pittsburgh, PA). The sequencing reads were mapped to the *M. acetivorans* C2A reference genome using breseq version 0.38.1 (35). Raw sequencing reads for DDN032, and the escape mutant are deposited in the Sequencing Reads Archive and will be made available upon publication.

### RNA extraction, sequencing, and transcriptomic analysis

Quintuplicate cultures of DDN032 and WWM60 were grown in 10 ml HS medium with 125 mM methanol and different concentrations of tetracycline as indicated in the text. 3 ml culture was removed for RNA extraction at an optical density between 0.2 and 0.6. The culture was immediately mixed 1:1 with RNA*later*, centrifuged at 10,000 x g for 10 minutes at 4°C, and the resulting pellet was applied to a Qiagen RNeasy Mini Kit (Qiagen, Hilden, Germany) and RNA extraction proceeded according to the manufacturer’s instructions. DNAse treatment, rRNA depletion, cDNA preparation and Illumina library preparation and sequencing were performed at SeqCenter (Pittsburgh, PA). Analysis of transcriptome data was carried out on the KBase bioinformatics platform (36). Briefly, raw reads were mapped to the *M. acetivorans* WWM60 genome using Bowtie2 (37), assembled using Cufflinks (38), and fold changes and significances values were calculated with DESeq2(39). Raw reads are deposited in the Sequencing Reads Archive (SRA) and and will be made available upon publication.

### Semi-quantitative western blotting

Late exponential-phase cultures were harvested by centrifugation, resuspended in 1 ml of lysis buffer (50 mM NaH_2_PO_4_, 2 U/ml Dnase I, 1 mM phenylmethylsulfonyl fluoride) and incubated at room temperature for ten minutes. An appropriate volume of a 5 M NaCl stock solution was added to bring the lysate to a final concentration of 300 mM NaCl. The lysate was then cleared by centrifugation (>10000 rcf, 4°C, 30 m, Sorvall Legend XTR (Thermofisher, Waltham, MA)) and the decanted supernatant was quantified with a microplate Bradford assay per manufacturer’s instructions (Sigma-Aldrich, Sant Louis, MO, USA).

Dilution series containing equal amounts of protein were prepared and separated on 12% Mini-PROTEAN TGX gels (BioRad, Hercules, CA, USA) by SDS-PAGE and then transferred onto 0.2 µM polyvinylidene difluoride (PVDF) membranes using the Trans-Blot Turbo system (BioRad) using Trans-Blot Turbo 0.2 PVDF transfer packs per manufacturer’s instructions. The membranes were then washed with phosphate-buffered saline containing 0.05% Tween20 (PBST) for five minutes at room temperature. Nonspecific binding was blocked by incubating in 5% milk in PBST for one hour at room temperature and four washes lasting five minutes each in PBST. The membranes were then incubated overnight at 4 °C in PBST with polyclonal rabbit antibodies raised against *mcrA* (1:10000 dilution) (GenScript, Piscataway, NJ, USA), washed four times for five minutes in PBST, and then incubated with anti-rabbit horseradish peroxidase (HRP) conjugate antibodies (1:20000 dilution) (Promega, Madison, WI, USA) for two hours at room temperature. Following four additional five-minute washes in PBST and three final washes in phosphate-buffered saline without Tween20, the membranes were developed with a five-minute incubation in Immobilon Western Chemiluminescent HRP substrate (EMD Millipore, Burlington, MA, USA) and imaged using a ChemiDoc XRS+ (BioRad). Sixty images were collected over one minute of imaging and the last images, which lacked oversaturation on any target bands, were selected for analysis using Image Lab.

## Author Contribution

G.L.C. contributed to conceptualization, data curation, formal analysis, supervision, methodology, and writing. G.A.D. contributed to data curation, formal analysis, methodology, and writing. D.D.N contributed to conceptualization, data curation, formal analysis, supervision, funding acquisition, project administration, methodology, and writing.

## Acknowledgements

We would like to members of the Nayak lab for their feedback and input on the manuscript.

## Data Availability Statement

All sequencing data have been deposited in the Sequencing Reads Archive and the bioproject number will be made available upon publication. All other data generated in this study will be made available upon request to the corresponding author.

## Supplementary Material

**Supplementary Table 1:**
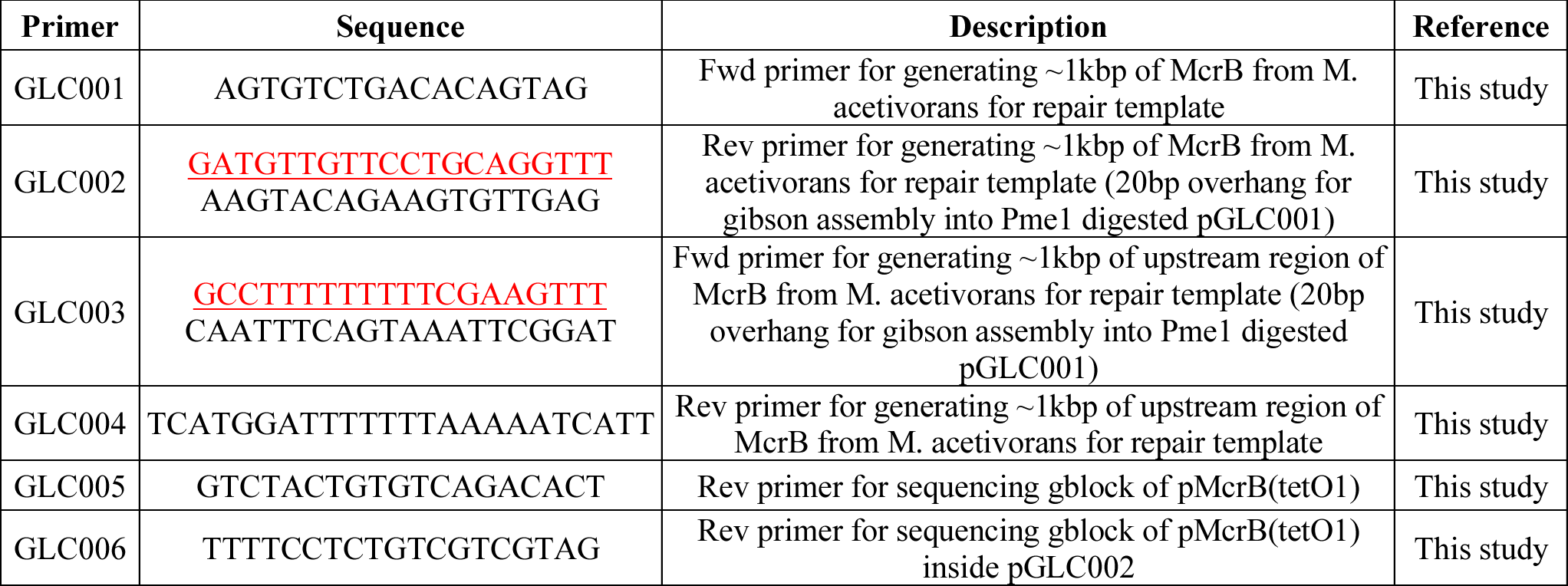
List of primers used in this study.

**Supplementary Table 2:**
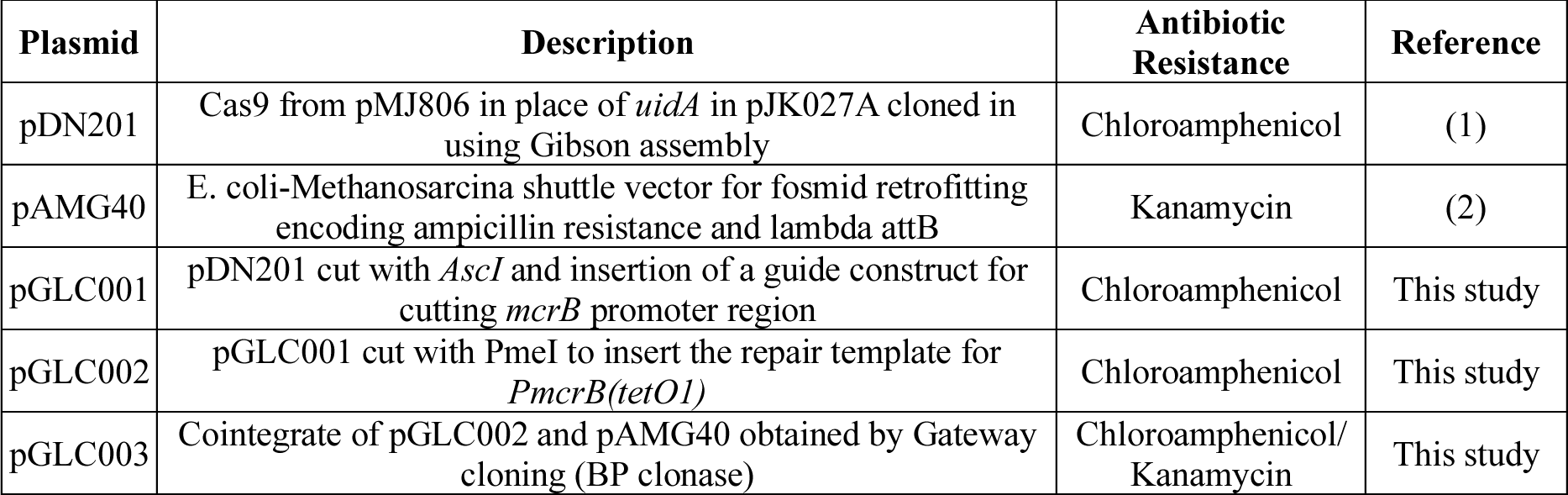
List of plasmids used in this study.

**Supplementary Table 3:** List of strains used in this study.

| Strain | Plasmid | Antibiotic Resistance | Genotype | Reference |
| --- | --- | --- | --- | --- |
| WM7959 | pDN201 | Chloroamphenicol | WM4489/pDN201 | (1) |
| WM3357 | pAMG40 | Kanamycin | WM1788/pAMG40 | (2) |
| DN112 | pGLC001 | Chloroamphenicol | WM4489/pGLC001 | This study |
| DN113 | pGLC002 | Chloroamphenicol | WM4489/pGLC002 | This study |
| DN114 | pGLC003 | Chloroamphenicol/<br>Kanamycin | WM4489/pGLC003 | This study |

**FIG S1:**
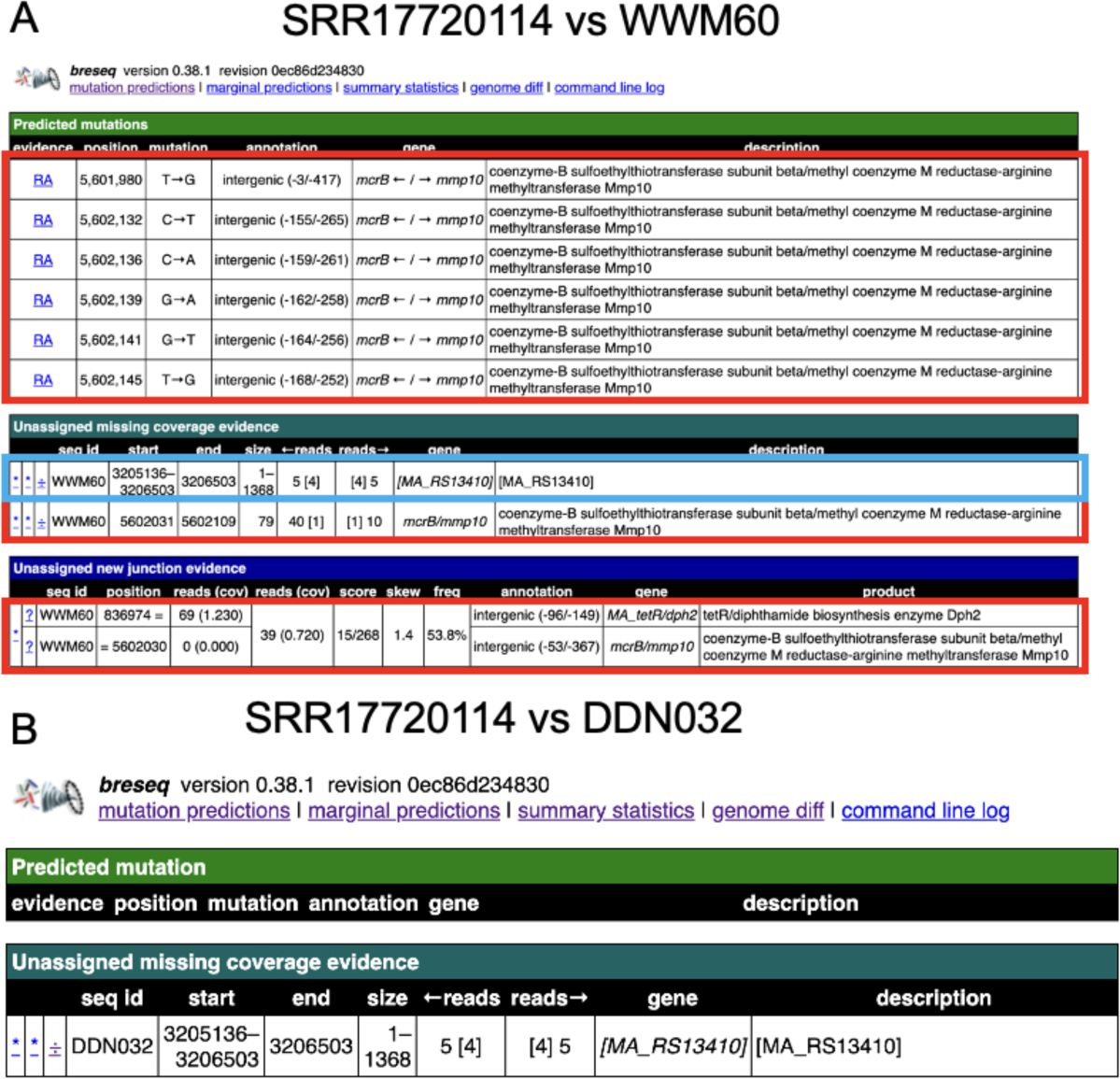
Genome resequencing. A) BreSeq analysis of DDN032 (SRR17720114) raw reads vs. WWM60. Changes in red correspond to expected modifications from inserting the *tetO1* operator site at the *mcr* promoter. Note: “Unassigned new junction evidence” is a spurious call due to the similarity between the native *mcr* promoter and the *mcr* promoter used to drive the t*etR* gene (MA_tetR). The missing coverage evidence for MA_RS13410 corresponds to a highly active transposase which has identical sequences spread throughout the genome often causing spurious missing coverage calls. B) BreSeq analysis of DDN032 (SRR17720114) raw reads vs. the expected full genome sequence of DDN032 shows no mutations and just the aforementioned low coverage of a transposase sequence.

**FIG S2:**
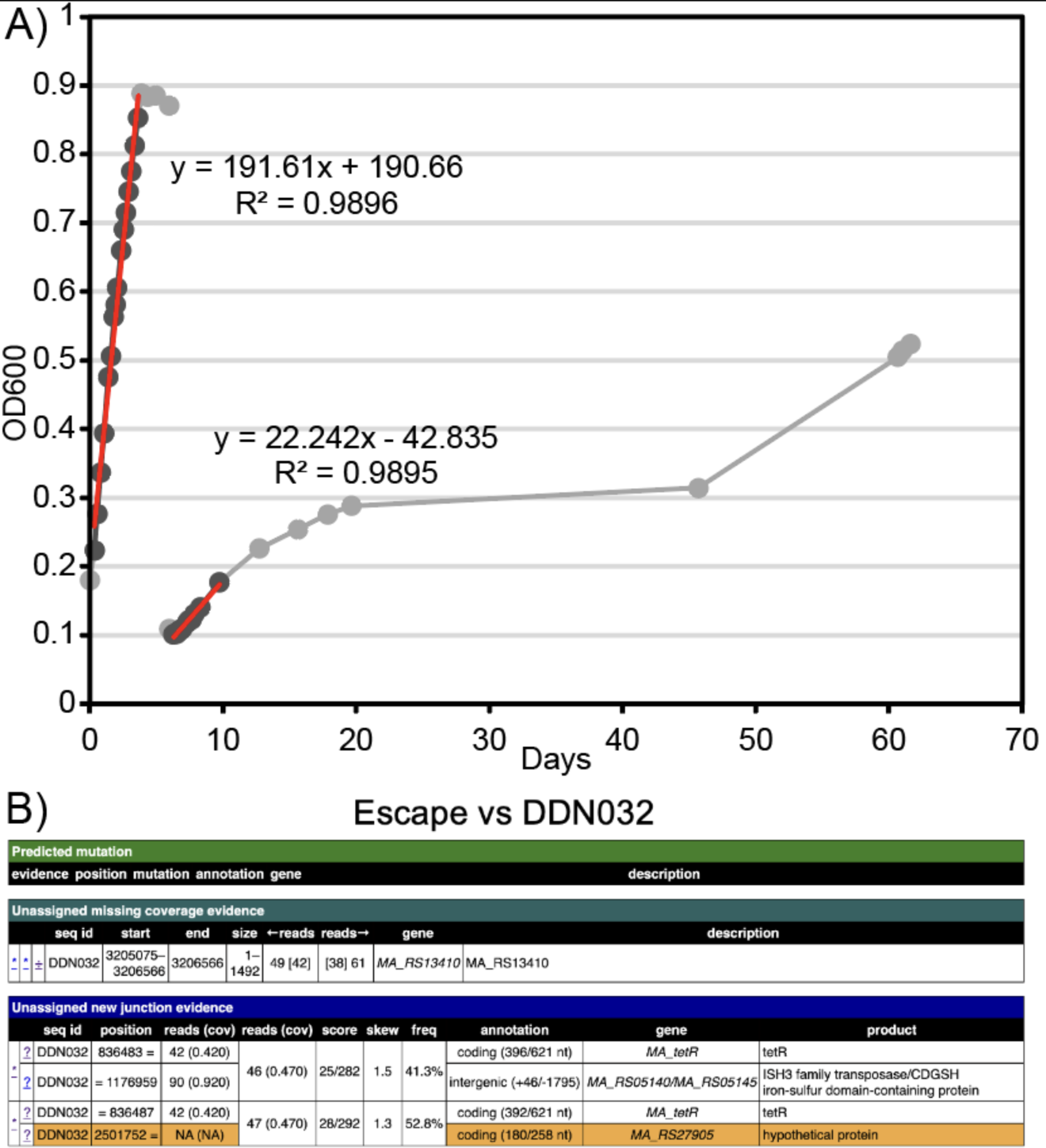
Long term incubation without tetracycline leads to escape mutants. A) Growth curves of DDN032 after washing and inoculation into tetracycline free media and subsequent passage. Slower linear growth is observed upon passaging consistent with **Fig. 1C**, which eventually levels off. Between 45 and 60 days, significant additional growth occurred, yielding a strain capable of exponential growth without the addition of tetracycline. B) resequencing of this escape mutant revealed the introduction of a transposon into the *tetR* gene.

**FIG S3:**
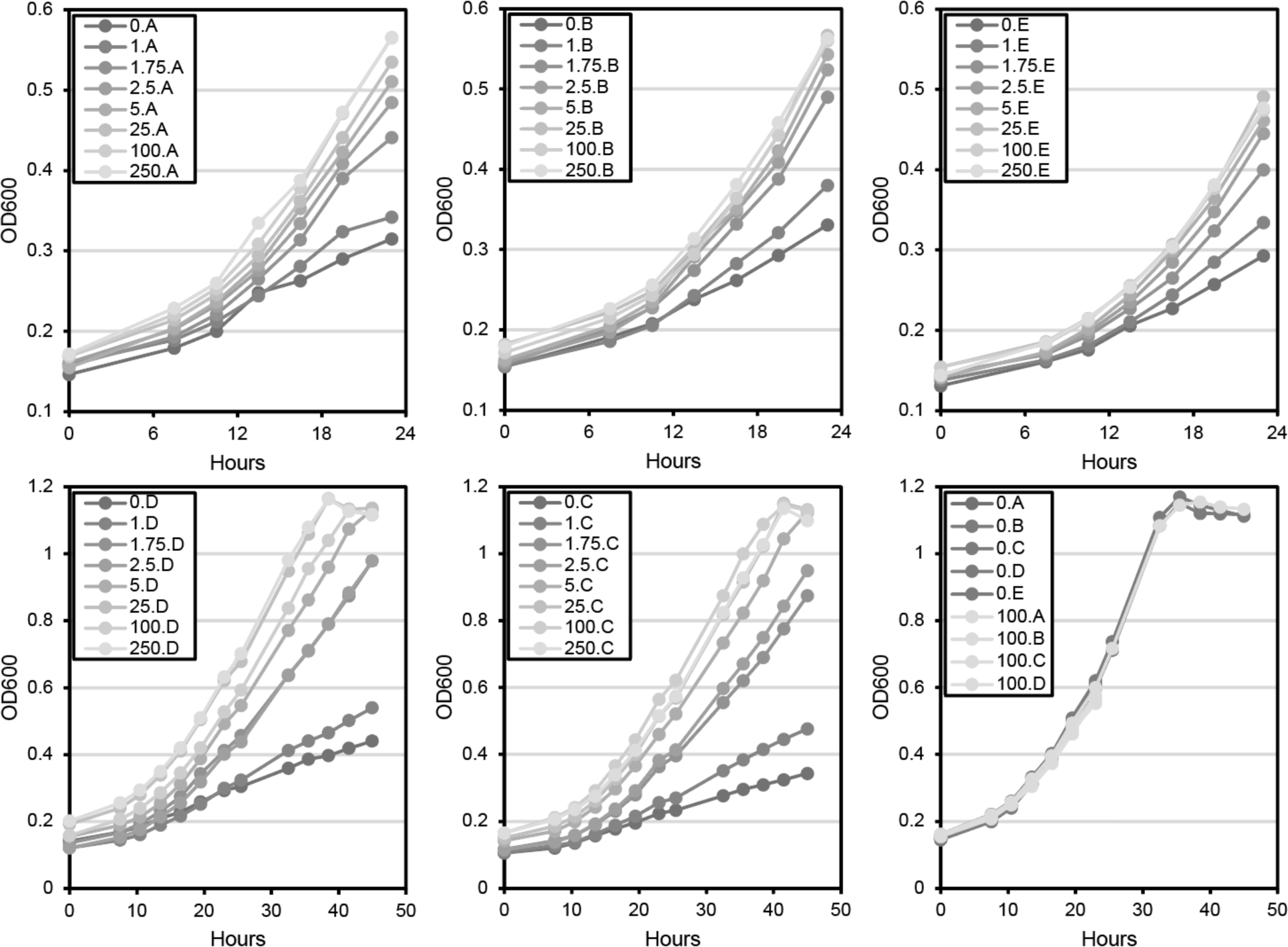
All growth curves from Experiment 1 shown in **Fig 2A-B**. The first five panels show replicates A-E of DDN032. Replicates A, B and E were harvested at 24 hours for RNA sequencing (**Fig 3-4**), while replicates C and D were allowed to continue growth. The final panel shows all growth curves of WWM60, three of which were also harvested at 24 hours. Legends show the concentration of tetracycline in µg/ml.

**FIG S4:**
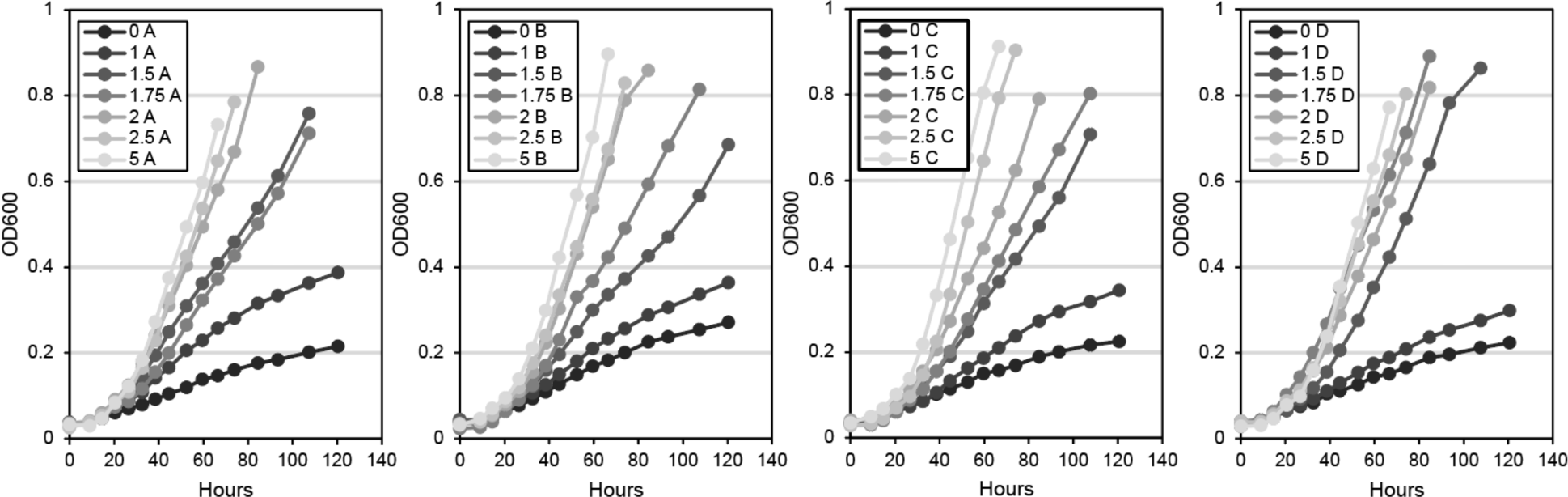
All growth curves from Experiment 2 shown in **Fig 2A-B**. Legends show the concentration of tetracycline in µg/ml.

**FIG S5:**
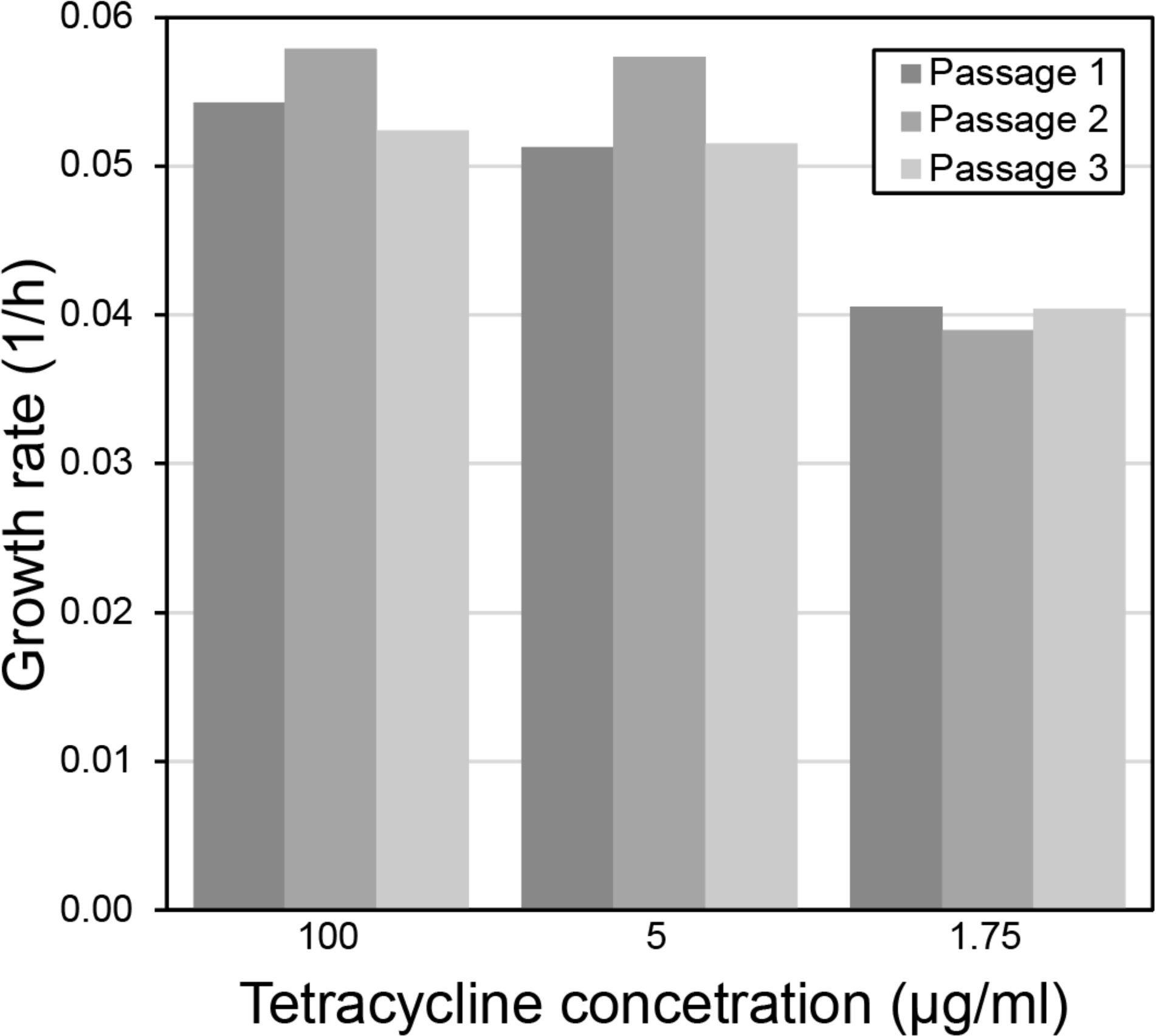
Doubling times for the exponential growth of sequential passages of DDN032 shown in **Fig 2C**.

**FIG S6:**
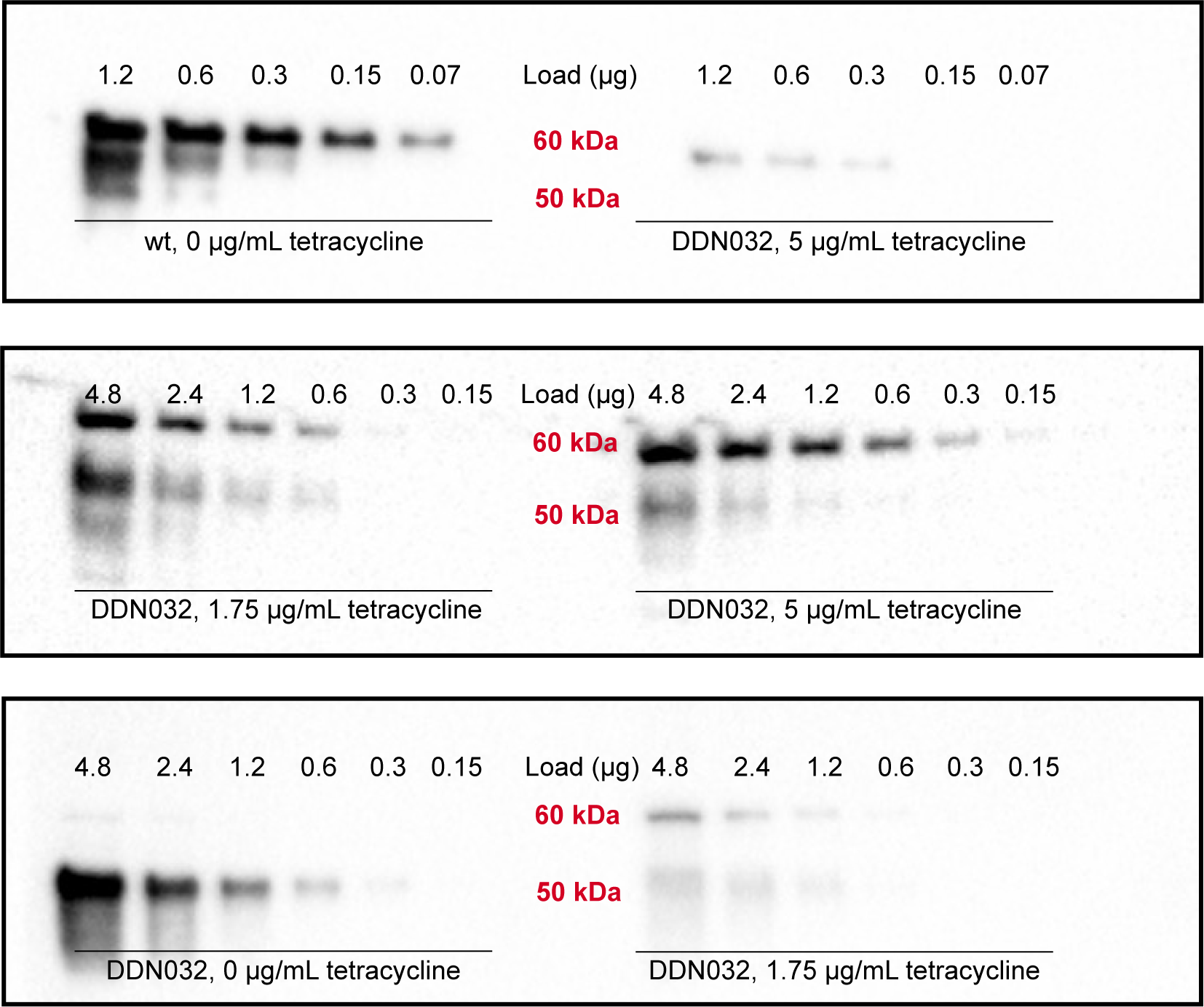
Representative full images of anti-McrA western blots. Comparisons between Mcr abundance at different levels of tetracycline concentrations (listed below each set of bands) should only be made with bands resulting from the same total protein load (µg) (listed above each band) on the same blot. Comparisons between blots is not possible. In addition to the 60 kDa band, associated with functional McrA, a secondary band at 50 kDa was detected. This band was confirmed by mass spectrometry to also be McrA and is presumed to be a degradation product, as has been observed in previous studies (3). The apparent high density of the secondary band in the 0 µg/mL tetracycline samples may be explained by the very low number of generations this culture completed before harvesting; these bands may represent the original pool of Mcr from the fully induced seed culture.

## Citations

1. Thauer RK. 2019. Methyl (Alkyl)-Coenzyme M Reductases: Nickel F-430-containing enzymes involved in anaerobic methane formation and in anaerobic oxidation of methane or of short chain alkanes. Biochemistry 58:5198–5220.

2. Culley DE, Kovacik WP, Brockman FJ, Zhang W. 2006. Optimization of RNA isolation from the archaebacterium *Methanosarcina barkeri* and validation for oligonucleotide microarray analysis. Journal of Microbiological Methods 67:36–43.

3. Ankel-Fuchs D, Jaenchen R, Gebhardt NA, Thauer RK. 1984. Functional relationship between protein-bound and free factor F430 in *Methanobacterium*. Arch Microbiol 139:332– 337.

4. Meyerdierks A, Kube M, Kostadinov I, Teeling H, Glöckner FO, Reinhardt R, Amann R. 2010. Metagenome and mRNA expression analyses of anaerobic methanotrophic archaea of the ANME-1 group. Environmental Microbiology 12:422–439.

5. Krukenberg V, Riedel D, Gruber-Vodicka HR, Buttigieg PL, Tegetmeyer HE, Boetius A, Wegener G. 2018. Gene expression and ultrastructure of meso- and thermophilic methanotrophic consortia. Environmental Microbiology 20:1651–1666.

6. Hallam SJ, Putnam N, Preston CM, Detter JC, Rokhsar D, Richardson PM, DeLong EF. 2004. Reverse Methanogenesis: Testing the hypothesis with environmental genomics. Science 305:1457–1462.

7. Krüger M, Meyerdierks A, Glöckner FO, Amann R, Widdel F, Kube M, Reinhardt R, Kahnt J, Böcher R, Thauer RK, Shima S. 2003. A conspicuous nickel protein in microbial mats that oxidize methane anaerobically. 6968. Nature 426:878–881.

8. Bonacker LG, Baudner S, Thauer RK. 1992. Differential expression of the two methyl-coenzyme M reductases in *Methanobacterium thermoautotrophicum* as determined immunochemically via isoenzyme-specific antisera. European Journal of Biochemistry 206:87–92.

9. Aldridge J, Carr S, Weber KA, Buan NR. 2021. Anaerobic production of isoprene by Engineered *Methanosarcina* species Archaea. Applied and Environmental Microbiology 87:e02417–20.

10. Chadwick GL, Joiner AMN, Ramesh S, Mitchell DA, Nayak DD. 2023. McrD binds asymmetrically to methyl-coenzyme M reductase improving active-site accessibility during assembly. Proc Natl Acad Sci USA 120:e2302815120.

11. Nayak DD, Liu A, Agrawal N, Rodriguez-Carerro R, Dong S-H, Mitchell DA, Nair SK, Metcalf WW. 2020. Functional interactions between posttranslationally modified amino acids of methyl-coenzyme M reductase in *Methanosarcina acetivorans*. PLOS Biology 18:e3000507.

12. Nayak DD, Mahanta N, Mitchell DA, Metcalf WW. 2017. Post-translational thioamidation of methyl-coenzyme M reductase, a key enzyme in methanogenic and methanotrophic Archaea. eLife 6:e29218.

13. Jin Q, Wu Q, Shapiro BM, McKernan SE. 2022. Limited mechanistic link between the Monod equation and methanogen growth: a perspective from metabolic modeling. Microbiology Spectrum 10:e02259–21.

14. Peterson JR, Labhsetwar P, Ellermeier JR, Kohler PRA, Jain A, Ha T, Metcalf WW, Luthey-Schulten Z. 2014. Towards a computational model of a methane producing Archaeum. Archaea 2014:e898453.

15. Cedervall PE, Dey M, Li X, Sarangi R, Hedman B, Ragsdale SW, Wilmot CM. 2011. Structural analysis of a Ni-Methyl species in methyl-coenzyme M reductase from *Methanothermobacter marburgensis*. J Am Chem Soc 133:5626–5628.

16. Aldrich HC, Beimborn DB, Bokranz M, Schönheit P. 1987. Immunocytochemical localization of methyl-coenzyme M reductase in *Methanobacterium thermoautotrophicum*. Arch Microbiol 147:190–194.

17. Jaenchen R, Gilles HH, Thauer RK. 1981. Inhibition of factor F430 synthesis by levulinic acid in Methanobacterium thermoautotrophicum. FEMS Microbiology Letters 12:167–170.

18. Moran JJ, House CH, Vrentas JM, Freeman KH. 2008. Methyl sulfide production by a novel carbon monoxide metabolism in *Methanosarcina acetivorans*. Applied and Environmental Microbiology 74:540–542.

19. Oelgeschläger E, Rother M. 2009. Influence of carbon monoxide on metabolite formation in *Methanosarcina acetivorans*. FEMS Microbiology Letters 292:254–260.

20. Rother M, Metcalf WW. 2004. Anaerobic growth of *Methanosarcina acetivorans* C2A on carbon monoxide: An unusual way of life for a methanogenic archaeon. Proceedings of the National Academy of Sciences 101:16929–16934.

21. Schöne C, Poehlein A, Jehmlich N, Adlung N, Daniel R, von Bergen M, Scheller S, Rother M. 2022. Deconstructing *Methanosarcina acetivorans* into an acetogenic archaeon. Proceedings of the National Academy of Sciences 119:e2113853119.

22. Shalvarjian KE, Nayak DD. 2021. Transcriptional regulation of methanogenic metabolism in archaea. Current Opinion in Microbiology 60:8–15.

23. Guss AM, Rother M, Zhang JK, Kulkkarni G, Metcalf WW. 2008. New methods for tightly regulated gene expression and highly efficient chromosomal integration of cloned genes for *Methanosarcina* species. Archaea 2:193–203.

24. Monod J. 1949. The Growth of bacterial cultures. Annual Review of Microbiology 3:371– 394.

25. Zheng K, Ngo PD, Owens VL, Yang X, Mansoorabadi SO. 2016. The biosynthetic pathway of coenzyme F430 in methanogenic and methanotrophic archaea. Science.

26. Deobald D, Adrian L, Schöne C, Rother M, Layer G. 2018. Identification of a unique Radical SAM methyltransferase required for the sp3-C-methylation of an arginine residue of methyl-coenzyme M reductase. 1. Sci Rep 8:7404.

27. Radle MI, Miller DV, Laremore TN, Booker SJ. 2019. Methanogenesis marker protein 10 (Mmp10) from *Methanosarcina acetivorans* is a radical S-adenosylmethionine methylase that unexpectedly requires cobalamin. Journal of Biological Chemistry 294:11712–11725.

28. Oelgeschläger E, Rother M. 2009. In vivo role of three fused corrinoid/methyl transfer proteins in *Methanosarcina acetivorans*. Molecular Microbiology 72:1260–1272.

29. Buan NR, Metcalf WW. 2010. Methanogenesis by *Methanosarcina acetivoran*s involves two structurally and functionally distinct classes of heterodisulfide reductase. Molecular Microbiology 75:843–853.

30. O’Donoghue P, Prat L, Kucklick M, Schäfer JG, Riedel K, Rinehart J, Söll D, Heinemann IU. 2014. Reducing the genetic code induces massive rearrangement of the proteome. Proceedings of the National Academy of Sciences 111:17206–17211.

31. Nayak DD, Metcalf WW. 2017. Cas9-mediated genome editing in the methanogenic archaeon *Methanosarcina acetivorans*. Proc Natl Acad Sci USA 114:2976–2981.

32. Eliot AC, Griffin BM, Thomas PM, Johannes TW, Kelleher NL, Zhao H, Metcalf WW. 2008. Cloning, Expression, and Biochemical Characterization of Streptomyces rubellomurinus Genes Required for Biosynthesis of Antimalarial Compound FR900098. Chemistry & Biology 15:765–770.

33. Metcalf WW, Zhang JK, Apolinario E, Sowers KR, Wolfe RS. 1997. A genetic system for Archaea of the genus *Methanosarcina*: Liposome-mediated transformation and construction of shuttle vectors. Proceedings of the National Academy of Sciences 94:2626–2631.

34. Sowers KR, Boone JE, Gunsalus RP. 1993. Disaggregation of *Methanosarcina* spp. and Growth as Single Cells at Elevated Osmolarity. Appl Environ Microbiol 59:3832–3839.

35. Deatherage DE, Barrick JE. 2014. Identification of mutations in laboratory-evolved microbes from next-generation sequencing data using breseq, p. 165–188. In Sun, L, Shou, W (eds.), Engineering and Analyzing Multicellular Systems: Methods and Protocols. Springer, New York, NY.

36. Arkin AP, Cottingham RW, Henry CS, Harris NL, Stevens RL, Maslov S, Dehal P, Ware D, Perez F, Canon S, Sneddon MW, Henderson ML, Riehl WJ, Murphy-Olson D, Chan SY, Kamimura RT, Kumari S, Drake MM, Brettin TS, Glass EM, Chivian D, Gunter D, Weston DJ, Allen BH, Baumohl J, Best AA, Bowen B, Brenner SE, Bun CC, Chandonia J-M, Chia J- M, Colasanti R, Conrad N, Davis JJ, Davison BH, DeJongh M, Devoid S, Dietrich E, Dubchak I, Edirisinghe JN, Fang G, Faria JP, Frybarger PM, Gerlach W, Gerstein M, Greiner A, Gurtowski J, Haun HL, He F, Jain R, Joachimiak MP, Keegan KP, Kondo S, Kumar V, Land ML, Meyer F, Mills M, Novichkov PS, Oh T, Olsen GJ, Olson R, Parrello B, Pasternak S, Pearson E, Poon SS, Price GA, Ramakrishnan S, Ranjan P, Ronald PC, Schatz MC, Seaver SMD, Shukla M, Sutormin RA, Syed MH, Thomason J, Tintle NL, Wang D, Xia F, Yoo H, Yoo S, Yu D. 2018. KBase: The United States Department of Energy systems biology Knowledgebase. 7. Nat Biotechnol 36:566–569.

37. Langmead B, Salzberg SL. 2012. Fast gapped-read alignment with Bowtie 2. 4. Nat Methods 9:357–359.

38. Trapnell C, Roberts A, Goff L, Pertea G, Kim D, Kelley DR, Pimentel H, Salzberg SL, Rinn JL, Pachter L. 2012. Differential gene and transcript expression analysis of RNA-seq experiments with TopHat and Cufflinks. 3. Nat Protoc 7:562–578.

39. Love MI, Huber W, Anders S. 2014. Moderated estimation of fold change and dispersion for RNA-seq data with DESeq2. Genome Biology 15:550.

## References

1. Nayak DD, Metcalf WW. 2017. Cas9-mediated genome editing in the methanogenic archaeon *Methanosarcina acetivorans*. Proc Natl Acad Sci USA 114:2976–2981.

2. Guss AM, Rother M, Zhang JK, Kulkarni G, Metcalf WW. 2008. New methods for tightly regulated gene expression and highly efficient chromosomal integration of cloned genes for *Methanosarcina* species. Archaea 2:193–203.

3. Aldrich HC, Beimborn DB, Bokranz M, Schönheit P. 1987. Immunocytochemical localization of methyl-coenzyme M reductase in *Methanobacterium thermoautotrophicum.* Arch Microbiol 147:190-194.

